# Evaluation of DNA conservation in Nile-Saharan environment, Missiminia, in Nubia: Tracking maternal lineage of “X-Group”

**DOI:** 10.1101/2020.04.02.021717

**Authors:** Yahia Mehdi Seddik Cherifi, Selma Amrani

## Abstract

**Objective:** We assessed DNA conservation using a range of archaeological skeletal samples from Sudan (Missiminia in Upper Nubia, 350 B.C.E to 1400 C.E) from the unfavorable conditions of the Saharan milieu and humidity of the Nile valley by tracking maternal lineage on the ‘X-Group’ (Ballaneans).

**Method:** We were able to extract, amplify, and sequence mt-DNA HVS-I (Sanger sequencing method) from 11 petrous bone samples, eight for the X-Group set and three for the reference set (one Christian, one Late Meroitic, and one Meroitic).

**Results:** It was possible to find the haplogroups (L1b, L2, L3, H2, N, T1a, X and W) and to carry out comparative data analysis in relation to haplogroup data cited in the literature. This investigation into the maternal lineage of X-Group (350 to 500 C.E.) origins allowed us to validate the efficiency of petrous bone sampling from ancient human remains from the Nile-Saharan milieu and established that the Ballaneans experienced an in-situ development with more admixture from the Levant region and North Africa.

**Conclusions:** Our study used mt-DNA (HVS-I) to look for the biological origins of the X-Group from Upper-Nubia and demonstrated the feasibility of ancient DNA research on skeletons from the Nile-Saharan environment. The use of Next Sequencing Generation (NGS) should optimize and improve the detection of shorter DNA strands and their sequencing in complete genomes from ancient skeletal remains (petrous bones) from hot and humid environments.

## 1. BACKGROUND

Archaeological samples, mostly bones and teeth, withstand the damaging outcomes of time and the environment and preserve the DNA within them. Recovering DNA from these surviving materials is influenced by numerous factors (Supplementary Figure 1S): experimental procedures [1–3], type of sample [4–6], taphonomy [6, 7], and climatic conditions [8–10]. However, not all elements having an impact on ancient DNA (aDNA) preservation and amplification are completely understood and are mainly dependent, as in previous observations [5], on environmental factors rather than on sample age [11, 12]. Here, we assessed DNA preservation using a range of archaeological skeletal samples from Sudan (Missiminia in Upper Nubia, 350 B.C.E. to 1400 C.E.).

### Ancient DNA studies in Nubia

The Nile valley of Sudanese Nubia is often considered as an incredibly rich and various archaeologically, although there are few investigations of aDNA in Nubia [13–15]. Themain aim of this study was to optimize the methodology and chances of successful sampling in Saharan environments. Paleogeneticists could address many important questions in the Nile valley, using aDNA analysis to seek the origin of the resident population and its evolution in relation to genetic and cultural admixture. Questions have been raised about the diversity of the population of Nubia because of its rich historical background, with its succession of three phases of Egyptian conquests, kingdoms and chiefdoms, and the constant pressure of desert tribes on the valley settlers.

While no proper anthropological storage facility or study was ever established in Nubia, most cranial materials were collected and transferred to the countries concerned with survey excavations: England, France, Denmark, and the United States [16–18]. Consequently, it is not unexpected that throughout the literature only three attempts to use aDNA from Sudanese Nubia have been published [13–15]. Fox (1997) [13] worked on Meroitic samples from Amir Abdellah (near Abri) using mitochondrial marker Hpa1 (np3,592) and suggesting sub-Saharan admixture. Francigny et al. (2013) [14] used teeth from four Nubian samples from different periods and archaeological contexts that succeeded in one amplification PCR (Christian); however, the typing autosomal STR is not confirmed. Sirak et al. (2014) [15] have worked on mt-DNA that highlighted only one sample (Christian), belonging to haplogroup L5, in favor of sub-Saharan admixture and confirming that aDNA could beanalyzed on samples from Saharan or sub-Saharan environments. Some articles were focused on the effect of military expeditions during the period of antiquity [19, 20] and others on genetic admixture pertaining to the Nubian corridor [21].

From ancient skeletal specimens, the success of DNA extraction and its reliability are complicated by degradation and contamination. For bone, local hydrology, pH, water content, oxygen levels, and soil composition have been identified as essential environmental parameters, contributing to the time period for the integrity of the material [22]. For DNA, temperature is considered an essential element before pH, soil content (salinity), and water content [23].

### Archaeological context

The Franco-Sudanese team, led by Andre Vila and the Khartoum National Museum, carried out five excavation campaigns in the years 1972 to 1975, enabling the archaeological sites to be surveyed along the Nile valley in Upper Nubia, along a length of 63 km, between the Dal cataract and the island of Nilwatti (Figure 1).

**Figure 1:**
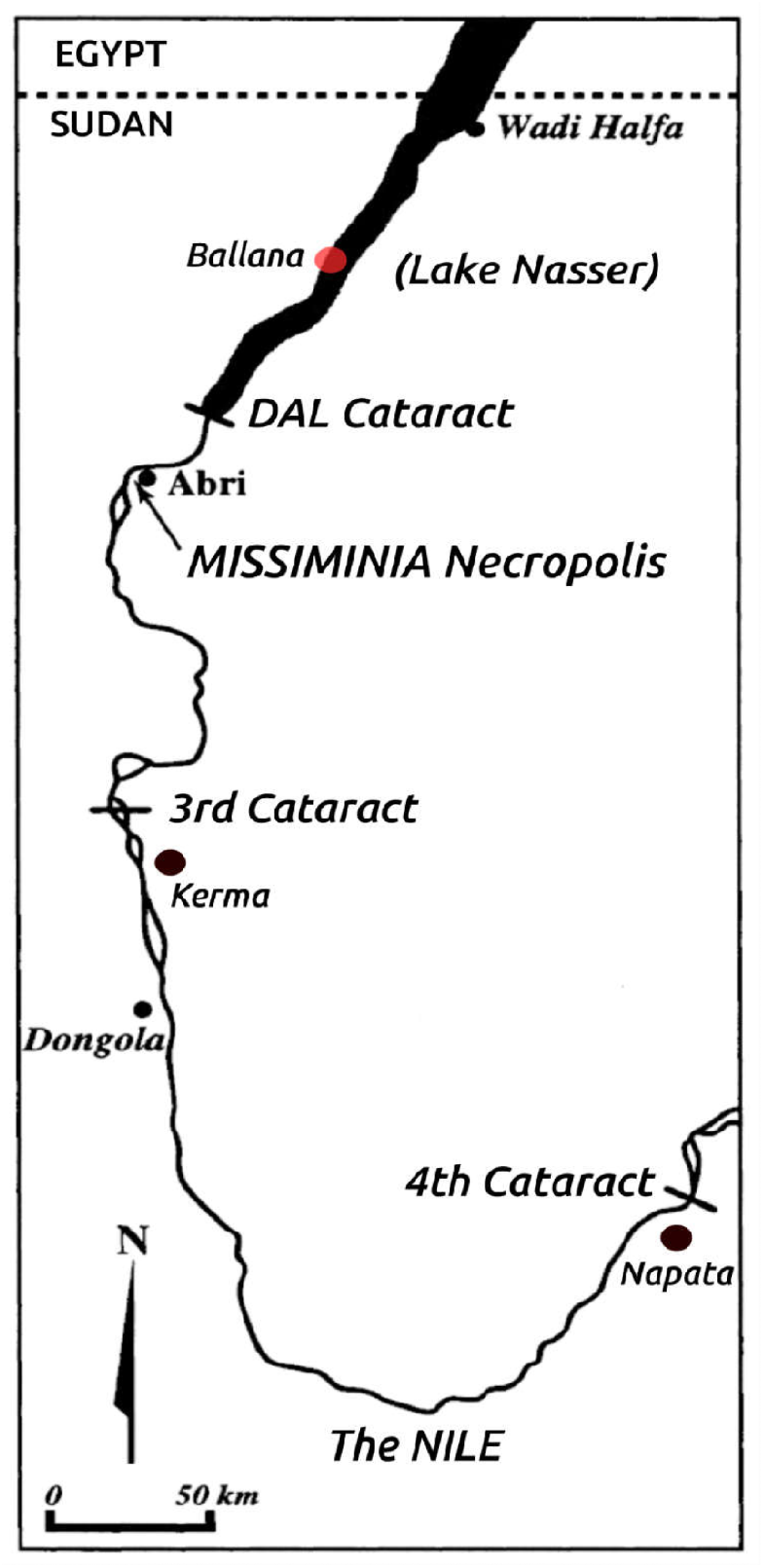
Missiminia necropolis, is located along the Nile valley in Upper Nubia (Sudan), from 200 km south of Wadi-Halfa, after the DAL cataract in the Abri district.

Important osteological material was collected and transferred to Limoges (France) for study in the laboratory during the year 1979. It consisted of 379 skulls and 346 mandibles, coming almost exclusively from the Missiminia Necropolis, which constitutes a funeral ensemble of 950 m in length by 100 m in width, about 700 meters south of the Abri’s souk (Figure 2). The major interest in this cranial collection is undoubtedly linked to the geographical situation of the necropolis in Sudanese Nubia, 200 km south of Wadi-Halfa (Figure 1). It is, in fact, the post-Pharaonic complex closest to the cultural centers of Napata and Meroe between the 3rd and 4th cataracts which were poor in terms of skeletal remains.

**Figure 2:**
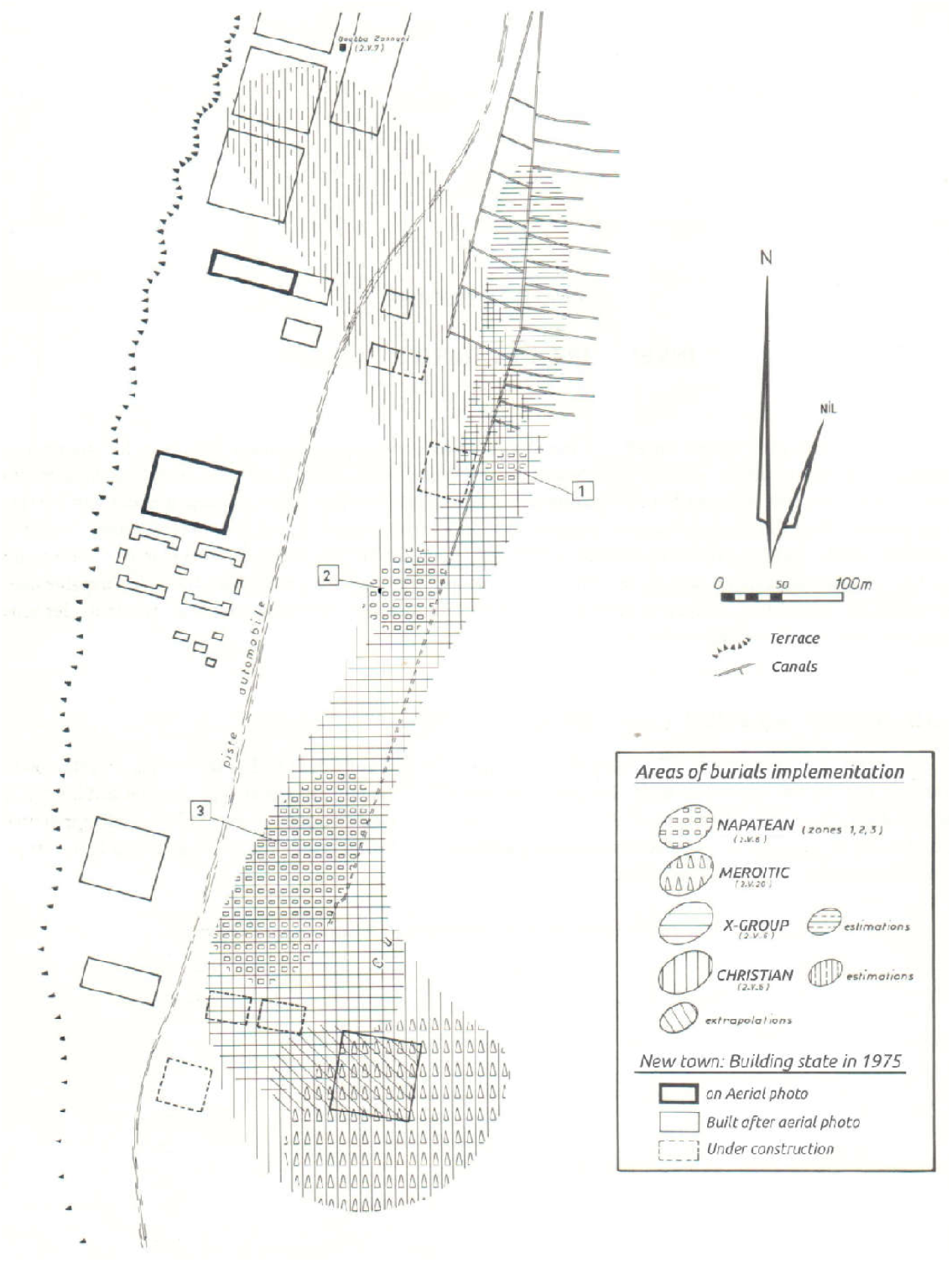
The areas of distribution of burials according to the different cultural phases and construction in progress of the new town of Abri-Missiminia (A.Vila of 1980).

At Missiminia, the proportion of skulls and mandibles in good macroscopic condition varied greatly, according to the cultural phase concerned: Napatean tombs with one subject out of 146 subjects (subjects from multiply used tombs, 16 pre-Meroitic samples studied came from other sites); Meroitic tombs and intrusive sepulchers (named “late Meroitic”) with 266 subjects out of 486 subjects; Ballanean Tombs (X-Group) with 56 subjects (54 skulls and 53 mandibles) out of 255 subjects from the graves of necropolis 2-V-6, except for one from 8-B-34 (cemetery of El Behel) near the Dal cataract; and Christian and undetermined graves with 91 subjects out of 269 subjects [24].

The causes of destruction are chiefly due to the action of moisture, depending on whether the tombs occupied the bed or, on the contrary, the banks of an old Nile arm, which also subjected them to exceptional floods. This is the case of the Napatean tombs and part of the Ballanean (X-Group) and Christian tombs, where the bones had become weak, if not powdered [17, 24].

Another cause is the plundering of the graves, causing the disintegration of the skeletons and breaking of the bones. But sometimes the upheaval and dispersal of the human remains resulted in a better conservation, as they were bathed in a sterile and desiccant environment: wind-borne sand. This was the case with the Meroitic tombs, used several times [24].

### About ‘X-Group’

The term X-Group Culture was used for lack of a more exact historical definition by George A. Reisner, who first discovered this civilization [25]. At Missiminia, the X-Group burial stretched through the length of the necropolis, on a strip of 300 by 80 meters (Figure 2). X-Group is known to be post-Meroitic, 350 to 550 C.E (Figure 3), associated with Ballanean Culture 300 to 600 C.E [17, 24, 26–28] due to their cultural changes, such as not using the Meroitic language, use of barrows in funeral practices, and ceramic innovation linked to Ballana [29]. These changes were accompanied by morphological variations [29, 30], and were based on an agrarian mode with a likely herding economy and isolated ‘Mediterranean break’ [31, 32]. Some included large barrows (between 5 and 20 m in diameter) which were to be those of persons who were to have, if not power, at least a certain social primacy. They can be considered true central sites [17, 29].

**Figure 3:**
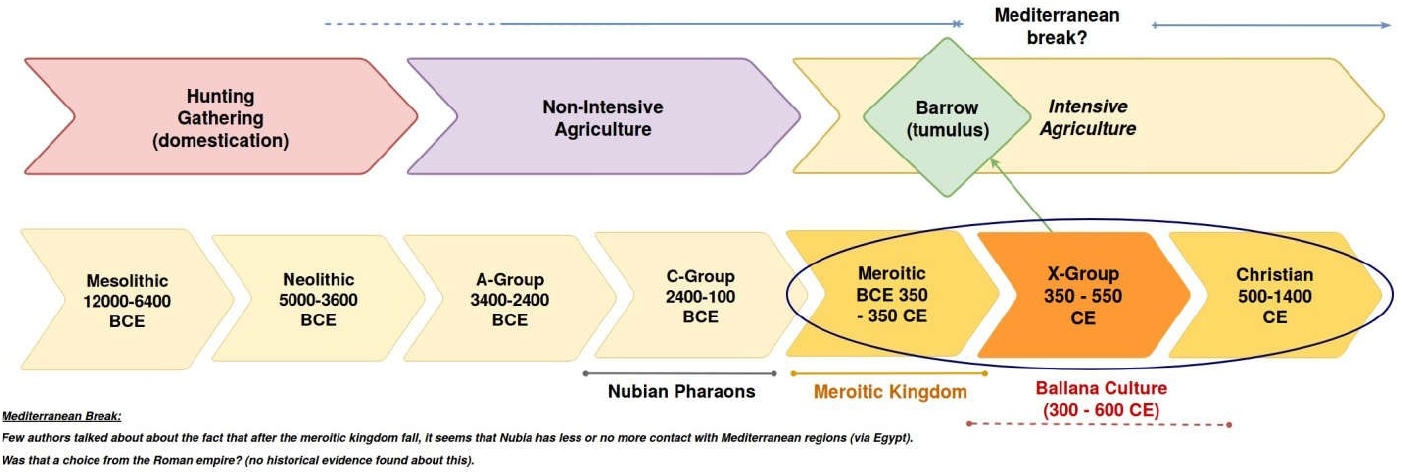
The Nubian population, had known seven specific periods of occupation or three broad phases spanning from the Mesolithic through the Christian era. Within the blue ellipse is our study area of interest, the main target being the X-Group.

The X-Group remains questionable and could be solved through aDNA. About their origins, there are two conflicting hypotheses or assumptions:

1. Outlier income: new people were associated with gene flow from both sub-Saharan and Mediterranean sources [17, 24, 29, 33–38].
2. In-situ: local people with genetic micro-evolution [13, 19, 20, 39]. A variation coming from a more open model of society is also proposed [15, 18, 29].

We conducted a study on 15 samples (petrous bones) from the Nubia, Missiminia necropolis to further investigate the following questions. These questions are essential for the optimized analysis of aDNA originating from a Nile-Saharan biotope:

- Is it possible to obtain a well-conserved aDNA from petrous bones from hot environments which are unfavorable for aDNA preservation?
- Are there differences between sampling on free and fixed petrous bone and the cranium [40]?
- Through maternal lineage determination, what is the history of the maternal part of the X-Group origins?

## 2. MATERIALS AND METHODS

All laboratory work was performed in the dedicated clean laboratory facilities at the Laboratory AMIS UMR 5288, University of Paul Sabatier — Toulouse III, according to strict aDNA standards [41, 42].

### Materials

The specimens reported in this study were skulls recovered during excavations at the Missiminia necropolis and studied osteologically. An exhaustive osteometric study was carried out by Billy, a brief odontological study by Verger-Pratoucy, and another brief paleopathological study by Dastugue [24], and a complete study was carried out on discrete cranial traits [17].

From the X-Group, 13 samples were selected and three other samples that occupied the Missiminia necropolis were added, one from each of the main groups (Table 1): Meroitic, Late Meroitic, and Christian (see ellipse in Figure 3). These served as the reference set to evaluate a difference in DNA preservation between these groups and the X-Group.

**Table 1.**
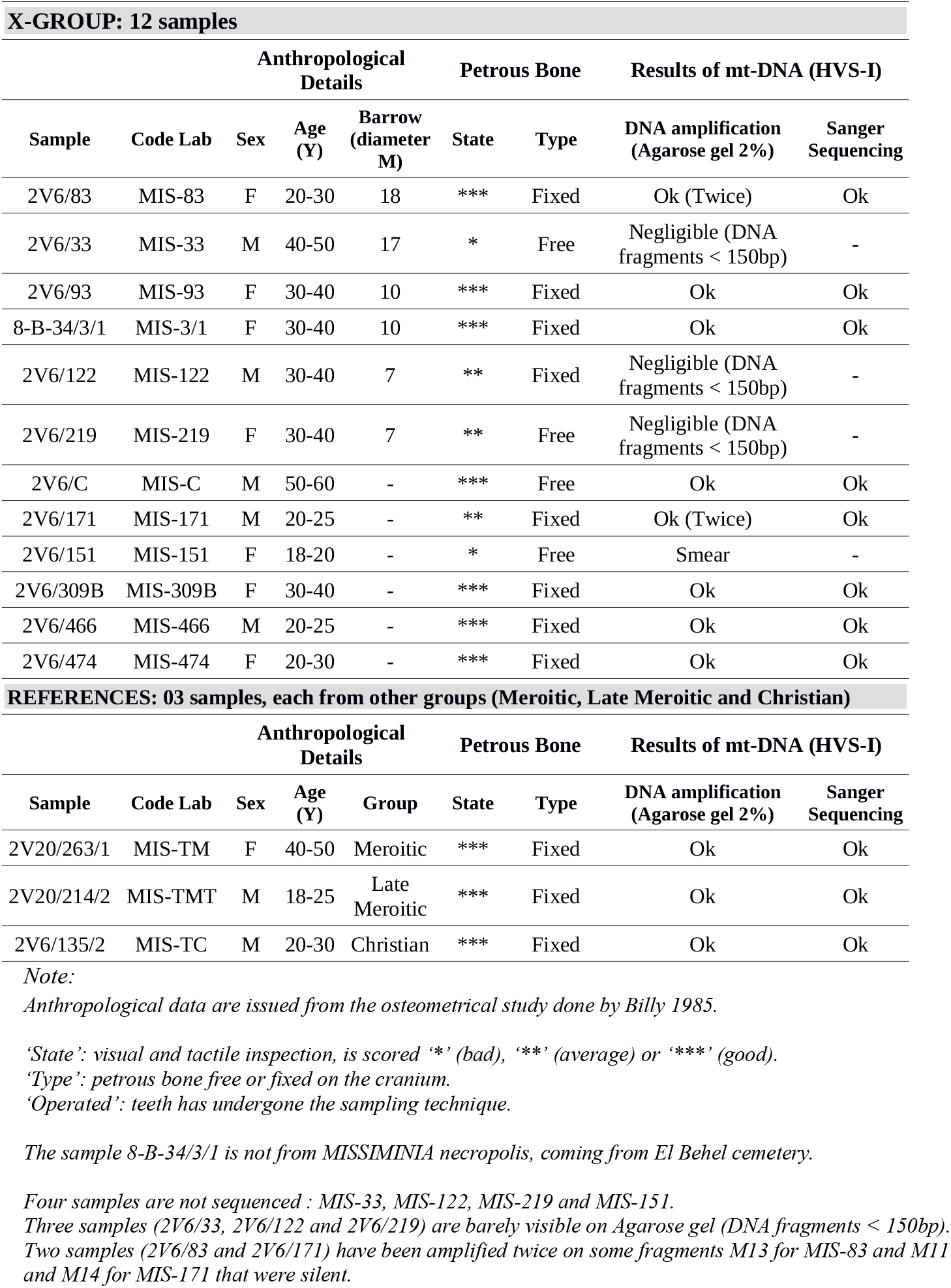
Samples details and results of mt-DNA (HVS-I) : 15 samples from Missiminia (X-Group and References)

Criteria for selecting the samples were primarily and consecutively: good specimens by visual and tactile inspection (scored as bad, average, or good), specimens having petrous bones (at least one), specimens (X-Group) found in a barrow (tumulus), and specimens, mainly X-Group, that could be related, or were at least reported as one set from previous studies [17]. Only four sample had a free petrous bone; the 12 others had the petrous bone fixed on the skull. Table 1 provides more details on the petrous samples.

### Methods

#### Authenticity criteria

Different precautions have been observed to reduce contamination with modern DNA. These include preserving the strict sterile conditions, minimizing the number of individuals handling the specimens, and determining the mitochondrial DNA polymorphism pattern of the researchers who had the direct contact with the specimens, as presented in Supplementary Table S1. Adherence to the best practice procedures for working with aDNA [43] was strictly maintained.

#### Sampling

To reduce the possibilities of contamination by modern DNA through handling, the sampling manipulations (cleaning and grinding) were performed by a single person wearing a full body suit dedicated to this task. The sampling of the specimens, previously irradiated by UV, was carried out in a specific room within a ductless fume hood. To further avoid contamination, all tools used were UV decontaminated within a Cross-Linker, and gloves were changed at key times.

The sampling protocol was adapted to the specimen. Thus, for free petrous bone or those originally fixed then detached using the Dremel (four cases), this included: pre-cleaning, abrasive cleaning of the surfaces before cutting (these two operations were carried out with a tool (Argofile MAXIMA PHP40 set, Japan), which can be fitted with forgers or saws depending on the needs), and harvesting into 5 ml tubes filled by a minimum of 300 mg of the bone powder [40]. The petrous bone attached to the skull was subjected to pre-cleaning (removing all contents of the cranial box), abrasive cleaning of the surfaces, drilling with Dremel 3000, and harvesting of the bone powder into 5 ml tubes filled by a minimum of 300 mg of the bone powder.

#### DNA operations

All aDNA manipulations were conducted in a dedicated aDNA facility with laminar flow hoods equipped with HEPA air filtration and inner UV light systems (one hood for DNA extraction, another for PCR setup).

##### Digestion

The lysis buffer, 555 *μ*l, containing 500 *μ*l of EDTA (0.45M), 50*μ*l of Proteinase K (2mg/ml) and of 5 *μ*l DTT (10mM), was added to a 2ml tube with 200-300 mg bone powder. Digestion was done overnight at 50°C with agitation (Specimix). Blank controls were used with each extraction series.

##### DNA Extraction

Any powder left was conserved into a plastic bag at −20^°^C and the supernatant was kept in the 5ml tube and 2.5 ml binding buffer was added. The mixture, 600 *μ*l, was added to a Qiagen Minelute Column, centrifuged 1 min at 8.000 rpm and the flow-through was discarded. This operation was repeated on the remaining mixture for each 600 *μ*l and finally was rinsed by 600 *μ*l wash buffer PE, centrifuged 1 min at 11.000 rpm, the flow-through was discarded, and the remainder was centrifuged 1 min again at 11.000 rpm to remove all the ethanol. DNA was eluted in 40 *μ*l Elution Buffer Kit Minelute Qiagen and 0.05% Tween, preheated at 37°C. It was incubated for 5 min at room temperature, and then centrifuged for 2 min at 13.000 rpm. Extracted DNA was stored at −20°C.

#### Amplification of DNA and mt-DNA Sequencing

##### Amplification of DNA

Considering the enormous capacity of PCR (Polymerase Chain Reaction) to amplify even a few copies of DNA sequences, modern DNA contamination has become a crucial problem. Another difficulty is the production of sufficient quantities of authentic DNA sequences to make a study conclusive, due to post-mortem DNA degradation processes (Supplementary Figure 1S) causing miscoding lesions potentially leading to sequence errors, thus raising the risk for preferential amplification of exogenous contaminant sequences.

To deal with this issue, we followed criteria authentication and adopted the use of four primer pairs, Mini-Primer set mt-DNA, to amplify the HVS-1 region in over lapping segments 146–165 bp in length as described in [44]. Blank controls were used to detect contamination, and positive controls (other ancient samples) were used to establish the effectivity of PCR set up in isolation from aDNA.

Amplification was carried out in a 25 *μ*l final volume PCR tube containing 23 *μ*l of the PCR mix (0.2 mM dNTP, 1.5 mM MgCl2, 0.2 mg/ml BSA, Buffer 1X (Gold ST*R 10X Buffer (Promega, Madison, WI)), primers at 600 nM and 2.5U AmpliTaq Gold DNA Polymerase (Applied Biosystems, Foster City, CA)) and 2 *μ*l of aDNA (for samples and positive controls) or water for PCR negative controls. After an initial 11 min at 95°C to denature DNA double strands, 40 cycles were done as follows: 95°C for 20 seconds, 50°C for 20 seconds, 72°C for 30 and final elongation step of 72°C for seven minutes. PCR products were visualized on a 2% Agarose gel.

##### Mt-DNA Sequencing

All PCR products, before sequencing, were purified using QIA quick PCR Purification Kit (Qiagen, Hilden, Germany) or were cleaned using a standard Exo-SAP protocol (5 *μ*l PCR product and 1 *μ*l of Exo-SAP, 30 min at 37°C / 20 min 80°C). Sequencing reactions were carried out on 96-well plates using BigDye Terminator v3.1 Cycle Sequencing Ready Reaction Kit (Applied Biosystems, foster City, CA) for 30 cycles, as follows: 95°C for 10 seconds, 50°C for five seconds, 60°C for two minutes. Sanger sequencing was run at Genopole Toulouse (France) on an ABI3730 Genetic Analyzer (Applied Biosystems).

#### Data analysis

DNA sequence analysis was accomplished by two different researchers, using the tools from NCBI BLAST (http://blast.ncbi.nlm.nih.gov/Blast.cgi) through alignment with the revised Cambridge Reference Sequence (rCRS) of mt-DNA (GenBank accession # NC 012920) to determine SNP differentiation.

All chromatograms were thoroughly inspected; using the SeqScanner Software (Applied Biosystems) by one researcher and the other used 4Peaks software (v1.8, A. Griekspoor and Tom Groothuis, http://nucleobytes.com/4peaks/index.html). Ambiguous base assignments were corrected manually. Alignment to the rCRS sequence was achieved using Bioedit software (v7.2.6, Tom Hall, http://www.mbio.ncsu.edu/BioEdit/bioedit.html). Haplogroups were defined using the haplogroup website (http://haplogrep.uibk.ac.at/) and SNP variations were referenced with the phylogenetic tree of global human mt-DNA variation (phylotree.org — mt-DNA tree Build 17 (18 Feb 2016)), to determine haplogroup assignment.

#### Haplogroup Affiliation

We collected a database of 3831 published HVS-I Haplogroups (with polymorphisms) from Arabia, North Africa, East Africa, West Africa, and the Near East that are described in Supplementary Table S2.

#### Statistical analysis

Statistical calculations and graphs were generated by R software (www.cran-project.org) using FactoMineR (factominer.free.fr) and factoextra (www.sthda.com/english/rpkgs/factoextra) packages for PCA and ggplot2 package (www.ggplot2.org) for graphics.

## 3. RESULTS

Mitochondrial HVS-I sequences were obtained for eleven specimens (73.3%) and can be classified into different haplotypes: African: L1, L2 and L3, Eurasian: N, H1, H2, N, T1, X and W (Table 1). All are still frequent in current East African, North African, Arab, and Near East populations (Supplementary Table S2).

### About Sampling

The X-Group’s 12 samples showed that four (36.36%) of the 11 specimens presenting a fixed petrous bone were labeled “average” state (Table 1), had their petrous bone detached, and had to be treated as free petrous bones. We found the protocol for skulls having a fixed petrous bone more constraining, as we were required to master the anatomical approach for drilling, temper contact with the bone to avoid overheating the powder (resulting in fragmentation or even destruction of the DNA contained within), and carry out a methodical cleaning of the crankcase and the drilling area to avoid any contamination of the powder.

### About DNA amplification

Four extractions of the total of 15 conducted (26.7%) were not conclusive and were not sequenced: three samples (MIS-33, MIS-122 and MIS-219 — Table 1) showed a negligible presence of DNA on Agarose gel (probably shorter fragments than our target 145-165 bp).

We were able to obtain DNA amplification in 73.3% of the cases, with only 4 failures: MIS-33, MIS-122, MIS-219, MIS-151 — Table 1. This failure was with 100% of the samples labeled “bad’” state (MIS-33 and MIS-151 — Table 1) and 66.7% of the samples labeled “average” state (MIS-122, MIS-219 — Table 1). We did not establish any notable difference in the success or failure of DNA extraction between the free or fixed petrous bone on the skull.

All the extraction and PCR blanks on the Agarose gel control were negatives, attesting to the absence of contamination during the procedures. We tried the procedure on two teeth samples, and failed to obtain an aDNA extraction and mitochondrial amplification.

### About Sanger Sequencing

All DNA amplifications obtained were successfully sequenced by the Sanger method, only MIS-151 (Table 1) had no conclusive sequencing (repeated twice without success). The references (MIS-TM, MIS-TMT and MIS-TC) were successfully sequenced whereas, on the X-Group’s 12 samples, eight were sequenced (66.7%) and analyzed with the assignment of a haplogroup (Table 1).

### General Diversity

The X-Group set showed some sharing of haplotypes: Four samples (50% of X-Group set) belonged to the African mega-haplogroup L with two individuals having haplogroup L3 (25% of X-Group set), one individual was labeled L2 and the other one L1. Regarding the Eurasian haplogroups, two individuals shared the X haplogroup (Table 2). The reference set showed one haplotype H2 (rCRS, no polymorphisms observed on the HVS-I) shared by two individuals (MIS-TM, MIS-TMT - Table 2).

**Table 2.** Haplogroups identification results (Alignment to the rCRS sequence) on HVS-I.

| **Code Lab** | **Range** | **Polymorphisms** | **Haplogroup** |
| --- | --- | --- | --- |
| MIS-83 | 16009-16391 | 16126C 16187T 16189C 16223T 16264T 16270T 16278T | **L1b** |
| MIS-93 | 16009-16391 | 16126C 16187T 16189C 16223T 16278T | **L2** |
| MIS-3/1 | 16009-16391 | 16182C 16183C 16189C 16223T 16278T | **X** |
| MIS-C | 16009-16391 | 16126C 16163G 16186T 16189C 16238T 16311C | **T1a** |
| MIS-171 | 16009-16391 | 16182C 16183C 16189C 16223T 16278T 16362C | **L3b** |
| MIS-309B | 16009-16391 | 16189C 16223T 16278T | **X** |
| MIS-466 | 16009-16391 | 16033T 16037G 16223T 16278T | **L3e** |
| MIS-474 | 16009-19391 | 16037G 16223T 16278T 16299T | **N** |
| MIS-TM | 16034-16385 | rCRS (no polymorphism) | **H2** |
| MIS-TMT | 16034-16365 | rCRS (no polymorphism) | **H2** |
| MIS-TC | 16024-16391 | 16037G 16223T 16292T 16295Y | **W1** |

### Haplogroup diversity

These haplogroups were compared with the database of current mt-DNA haplogroups collected from the literature (Supplementary Table S2). The distribution among the X-Group seemed to be highly diverse, belonging to common Eurasian (N, T1, X) and sub-Saharan haplogroups (L1–L3), as expected. However, they were less diverse in the reference set; surprisingly they belonged only to common Eurasian haplogroups (H2 and W1).

### Mitochondrial Diversity

Due to the low number of aDNA sequences, and in order to observe the mt-DNA patterns of the X-Group population within a broad geographical context, we performed a Principal Component Analysis (PCA) based on the frequencies (Table 3) from the literature cited (Supplementary Table S2)(Figure 4).

**Table 3.** Haplogroup frequencies (%) (based on groups of populations), modern (n= 3831) populations used in the population genetic analyses.

| **Population** | **Code** | **n** | **L1b** | **L2** | **L3b** | **L3e** | **N** | **T1a** | **X** | **H2** | **W1** | **Others** |
| --- | --- | --- | --- | --- | --- | --- | --- | --- | --- | --- | --- | --- |
| Arabia | ARA | 365 | **1.37** | **3.56** | **1.64** | **1.10** | **25.48** | **10.68** | **9.59** | **8.77** | **19.45** | 18.36 |
| Near-East | NEE | 628 | **0.16** | **0.64** | **1.11** | **0.96** | **17.36** | **11.46** | **10.67** | **9.71** | **16.24** | 31.69 |
| North-Africa | NAF | 1676 | **4.18** | **9.01** | **3.58** | **5.37** | **1.79** | **1.19** | **8.95** | **24.46** | **9.25** | 32.22 |
| East-Africa | EAF | 969 | **1.75** | **16.31** | **20.74** | **26.83** | **0.52** | **1.03** | **1.03** | **3.41** | **0.31** | 29.41 |
| West-Africa | WAF | 193 | **11.40** | **51.81** | **8.29** | **10.88** | **0.00** | **0.00** | **0.00** | **0.00** | **0.00** | 17.62 |
| **Total** |  | **3831** | **3.00** | **11.12** | **7.57** | **7.83** | **5.48** | **3.21** | **5.66** | **13.99** | **8.64** | 33.49 |
| **Missiminia**<br>**X-Group only** | **MIS** | **8** | **12.50** | **12.50** | **12.50** | **12.50** | **12.50** | **12.50** | **25.00** | **0.00** | **0.00** | 0.00 |

*Note:*
*Litterature references are listed in Supplementary Table S2 (sources with literature cited details).*
*‘Others’ could include sub-clads from L1, L2 or L3, H1, and also from other haplogroups that were not listed within the X-Group (L0, L4, M, I, HV, ...).*

**Figure 4:**
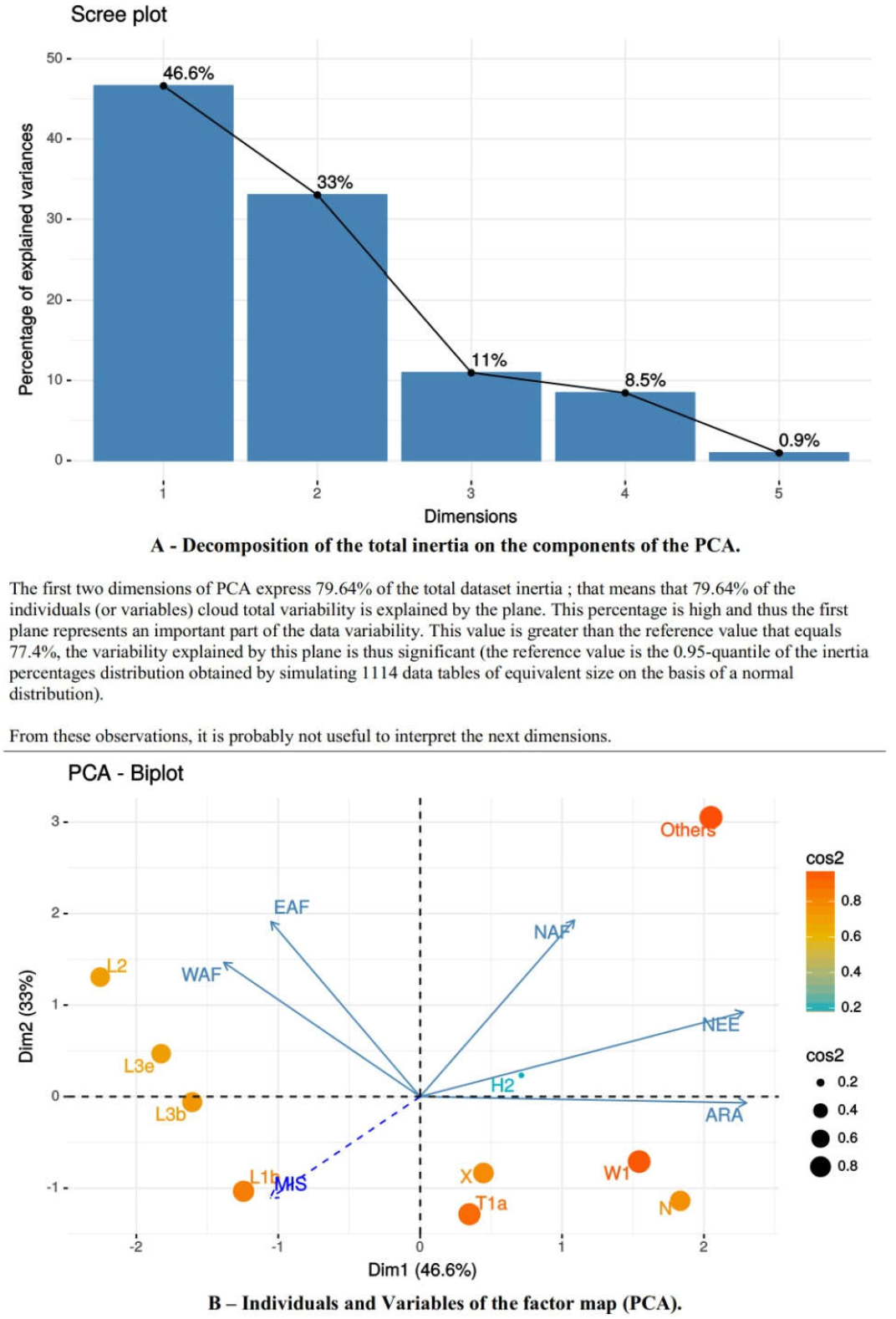
This PCA resumes the Haplogroups diversity and distribution (Table 3) compared to the database collected in literature (Supplementary Table S2).

Principal components 1 and 2 (Dims 1 and 2 in Fig. 4) account for 79.6% of the total variation (46.6% and 33%, respectively). The two-dimensional pattern displays distinct population clusters: the sub-Saharan populations (East African EAF and West African WAF) located in the most negative portion of axis 1; the Levant region populations (Arabian-ARA and Near Eastern-NEE) in the upper-right quadrant, respectively; and North Africans (NAF) sandwiched between the sub-Saharan cluster and Levant Region populations, but closer to the latter. If we detail the position of our X-Group (MIS) samples in the overall mitochondrial diversity, the sub-Saharan (EAF and WAF) are most closely related to them whereas those from the Levant region (ARA and NEE) are the furthest away. The North Africans (NAF) are the complete opposite to the X-Group.

The population positions on the PCA Biplot (Figure 4) are explained by the distinction between African and Eurasian haplogroups. As expected, sub-Saharan populations are characterized by high frequencies of L lineages (L1b, L2, L3b and L3e) whereas Eurasians are mainly associated with lineages H2, N1, T1a, X and W. Then, East Africans show high frequencies of L3 (L3b and L3e) and L2 lineages, and Near Easterners are mainlyassociated with lineages H2, X, and N1. Finally, the central position of North Africans is explained by higher frequencies of Eurasian lineages with respect to the African haplogroups, the small frequencies of Eurasian and X lineages, and the presence of the H2 haplogroup.

Overall, the PCA biplot (Figure 4) based on mitochondrial haplogroup frequencies reflects a clear differentiation between the X-Group and non-sub-Saharan groups (ARA, NEE and NAF) but a close relationship between the X-Group and sub-Saharan populations (EAF and WAF).

## 4. DISCUSSION

The purpose of this study was to assess DNA preservation using a range of archaeological skeletal samples from Sudan (Missiminia in Upper Nubia), from the unfavorable conditions of the Saharan biotope, by tracking the X-Group. Here, we discuss aDNA quality, sequencing, and maternal lineage through mt-DNA haplogroups.

### Ancient DNA preservation and its quality

In addition to precautions carried out to prevent contamination, the monitoring and the control of DNA presence and its quality allowed for increasing compliance with the technical procedures adapted to carry out this study.

In our case, we noticed that samples labeled “bad/average” state (visual) corresponded to low DNA presence on Agarose gel, and also to the failure of amplification and/or sequencing procedures (Table 1), as reported by [12]. The control of 2% Agarose gel prior tosequencing revealed that, the extraction and PCR blanks were free of any contamination. This was confirmed by the absence of DNA after sequencing. The post-treatment by the Sanger sequencing method, by their length and readability (too many N or non-homogeneous curves and not individualized), allowed us to detect miscoding lesions and make decisions on running another sequencing for the relevant samples.

The non-negligible role of environmental parameters in the diagenetic processes undergone by DNA in skeletal remains [45], and both mold and soil environments favorable to the proliferation of microorganisms, were very detrimental to bone DNA conservation because more than 97% of the bone DNA was lost [46]. On the other hand, the extraction method used has been demonstrated to remove on average 64% of the microbial DNA from bone powder but only 37% of the endogenous DNA (from the organism under study), increasing the percentage of informative sequences by a factor of two on average [47], and could explain our good success rate for viable aDNA extraction from archaeological materials from this region.

### Sequencing

Haplogroup affiliations and HVS-I sequences were determined for 11 subjects from Missiminia necropolis, only eight subjects were identified being from the X-Group. Two subjects (MIS-3/1 and MIS-309B — Table 2) were affiliated with the X haplogroup, even if a researcher (AMIS-R03 — Supplementary Table S1) on the sub-Haplogroup X1’3 caused a possible contamination which is not excluded without the results of the HVS-II region which may or may not invalidate this possibility. This would require using Next Generation Sequencing (NGS) to distinguish between ancient and modern DNA. On these two subjects, we should reserve discussion of the results.

Three subjects (MIS-33, MIS-122 and MIS-219. Table 1) showed a negligible presence of DNA on Agarose gel control, an indication of the availability of DNA fragments below our target of 145-165 bp [44]. This aDNA could possibly be obtained through NGS technology, more suited for short fragments, 40 to 90 bp [48, 49].

### mt-DNA Haplotyping (Maternal Lineage)

By identifying the affiliation of the haplogroups and their comparison with the cumulative data from the literature (Supplementary Table 1), we can discuss each haplogroup to verify hypotheses on the X-Group origins. We found an influx of sub-Saharan African ancestry after the Meroitic Period, which corroborates the findings of Schuenemann et al. (2017) [50]. L1b and L2 haplogroups are mostly present in West Africa, respectively at 11.4% and 51.8% [51], and also present in North Africa [52–55]. The presence of L1b in the X-Group seems to be due to genetic flow from West to East Africa probably via the nomadic tribes, Berbers in northern Africa [52, 53] and Tuareg in the Sahara [55]. While the presence of L2, in the X-Group, seems to be part of its equivalent presence in West and East Africa with shared genetic matches with North Africa [56].

L3b and L3e haplogroups are mostly present in East Africa, respectively at 20.7% and 26.8%, mainly in Sudan [57] and in Ethiopia [58, 59], thus, that they are within the X-Group is possible evidence of this in situ evolution, comprising the possibility of a southsouth exchange for L3b or an east-west exchange for L3e [56, 60–62].

N, T1a and X haplogroups are mostly present in Arabia and the Near East, respectively at 25.5%, 11.5%, and 10.0%, and still present in North Africa at less than 1.8% [63]. Without minimizing the question of probable unrecognized contamination (Researcher AMIS-R03 — Supplementary Table 1), the presence of this haplogroup X, like T1a and N haplogroups in the X-Group, suggests the possible exchange with North Africa and the Levant region (Near East).

Haplogroups W1 and H2 were not found in the X-Group, just in the reference set respectively still present at around 18% in Arabia and the Near East and almost 25% in North Africa [52]. The presence of this haplogroup in the Christian reference (MIS-TC) qualified as Asian, and could suggest by its confirmed presence in Ethiopia [59] an in-situ sub-Saharan exchange. As for the X haplogroup, the presence of this haplogroup H (H2 — rCRS) within the reference set (MIS-TM, MIS-TMT — Table 2) corroborates the possible exchange with North Africa and the Levant region (Near East). This also suggests a possible genetic continuity between the Meroitic (MIS-TM) and Late Meroitic (MIS-TMT), which some authors [17, 18] had proposed from an anthropological point of view, as opposed to the archaeological view point of these two sets uniting under the common name of Meroitic.

About the PCA Biplot (Figure 4): Could the significant negative correlation between the X-Group and the North Africans (NAF) possibly be a Mediterranean Break effect on the X-Group? We could not make any clear statement about that as we have a small number of samples and aDNA sequences.

This study on 15 samples (petrous bones) from Nubia, Missiminia necropolis (350 B.C.E to 1400 C.E.), leads us to verify an essential question about the optimal analysis of aDNA on samples from the Nile-Saharan environment. Although the Saharan climate is detrimental to the preservation of DNA, and whilst it is associated with the disadvantages of moisture and salinity (on bone and DNA) that could be added by proximity to the Nile in our study, we were able to extract aDNA (mt-DNA), amplify it and then sequence it successfully on 11 samples (73.3% success rate), and track the maternal lineage of the X-Group. In general, we observed that “bad/average” visual state correlates to failure in biomolecular procedures, as reported in the literature [12]. However, the study lead us to rethink and optimize new perspectives such as the use of NGS technology to access shorter fragments (40 to 90 bp) while reconstructing whole genomes (mt-DNA, autosomal and Y-chromosomes) that allow new interpretations, integrating the measurement of the quality of the DNA with a bio-analyzer and the massive sequencing at high bit rate, genotyping, validating the presence of modern DNA, the quality of the old DNA obtained, and rebuild phylogenies and phylogeographies based on notions of kinship or even on regional movements of populations. We answered the question of X-Group origins and validated the thesis of an in situ micro-evolution from African local-south and local-east exchanges with more admixture and a more open society than expected towards the Levant region (Near/Middle East) and North Africa as found on haplogroups analysis. However, a more detailed study on a larger sample would be required to assess these statements based on sampling subjects from 300 specimens, with 36 of X-Group and 27 of the pre-Meroitic period found in our collection.

## 5. CONCLUSION

Our tracking of the X-Group of the necropolis of Missiminia (Upper Nubia — Sudan, 350 B.C.E to 1400 C.E) via the mt-DNA haplogroups (Sanger sequencing method) allowed us to test hypotheses about their origins, to validate the feasibility of an aDNA study on petrous bones from a Saharan climate with a humid contribution via the Nile (the most unfavorable hot and humid milieu for DNA preservation), and to confirm endogenous DNA protected from climatic variances within the petrous bone.

Our analysis of 15 samples (petrous bones) from the Missiminia necropolis, answered our questions:

- Yes, it is possible to obtain well conserved aDNA on petrous bones from hot and humid environments (we obtained a 73.3% success rate).
- No, there is no notable difference between free and fixed petrous bone. Moreover, we do not demonstrate the significance of such a difference. (This could also be done within another experimental design).
- The maternal part of the X-Group origins history suggested a more diverse model society than expected, based on Sub-Saharan with Eurasiatic admixture, from Levant and North Africa (to be confirmed by a further study including more samples and a different experimental design).

Our study shows that paleogenetic studies on around 2000-year-old skeletons, from Saharan-type environments (even when coupled with the high humidity of the Nile), are feasible on petrous bones free or fixed to the skull. Strict rules to avoid contamination and the conduct and follow-up of technical procedures for mt-DNA via the Sanger method sequencing. The transition to Next Generation Sequencing (NGS) technology should improve the detection of shorter DNA strands and their sequencing in complete genomes on ancient skeletal remains from hot and humid environments.

## 6. ACKNOWLEDGEMENTS

We are very grateful to the AMIS lab team, Toulouse, France for their helpful discussion and comments during the preparation of the manuscript, and to Ryan Schmidt (Ron Pinhasi Lab, UCD, and Ireland) for his helpful discussion and comments about bioinformatics and methodology on the Sanger method and analysis. The present version greatly benefited from the comments provided by the Editors and two anonymous reviewers.

## 7. AUTHOR CONTRIBUTIONS

Conceptualization: Yahia Mehdi Seddik CHERIFI.

Formal analysis: Yahia Mehdi Seddik CHERIFI, Selma AMRANI.

Investigation: Yahia Mehdi Seddik CHERIFI.

Methodology: Yahia Mehdi Seddik CHERIFI, Selma AMRANI. Visualization: Yahia Mehdi Seddik CHERIFI, Selma AMRANI.

Writing — original draft: Yahia Mehdi Seddik CHERIFI, Selma AMRANI.

Writing — review & editing: Yahia Mehdi Seddik CHERIFI.

## 8. CONFLICT OF INTEREST

The authors have none to declare.

