## Supplementary material for "Evaluation of DNA conservation in Nile-Saharan environment, Missiminia, in Nubia: Tracking maternal lineage of “X-Group”": Supplemtary Figure S1

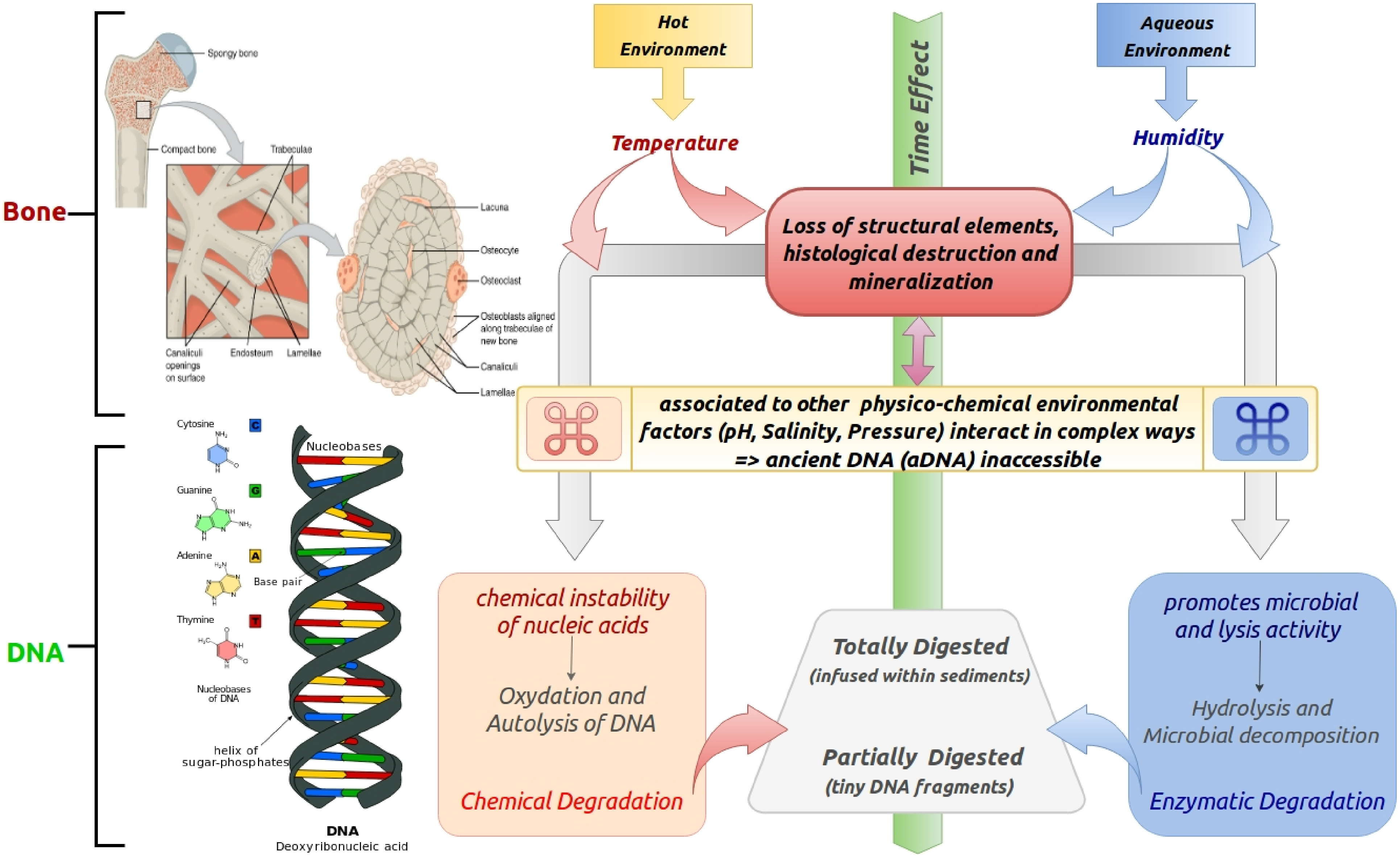
Bone
Spongy bone
Compact bone
Trabeculae
Canaliculi openings on surface
Endosteum
Lamellae
Lacuna
Osteocyte
Osteoclast
Osteoblasts aligned along trabeculae of new bone
Canaliculi
Lamellae
DNA
Cytosine
C
Guanine
G
Adenine
A
Thymine
T
Nucleobases of DNA
Nucleobases
Base pair
helix of sugar-phosphates
DNA
Deoxyribonucleic acid
Hot Environment
Temperature
Time Effect
Aqueous Environment
Humidity
Loss of structural elements, histological destruction and mineralization
associated to other physico-chemical environmental factors (pH, Salinity, Pressure) interact in complex ways
=> ancient DNA (aDNA) inaccessible
chemical instability of nucleic acids
Oxydation and Autolysis of DNA
Chemical Degradation
Totally Digested
(infused within sediments)
Partially Digested
(tiny DNA fragments)
promotes microbial and lysis activity
Hydrolysis and Microbial decomposition
Enzymatic Degradation

Table S1

Personal laboratory (worked on samples) HVS-I and HVS-II mitochondrial sequences

| **Researcher Id** | **Range** | **Polymorphisms** | **Haplogroup** |
| --- | --- | --- | --- |
| AMIS-R01 | 16012-263 | 16304 16311 146 263 | **H5a5** |
| AMIS-R02 | 16012-263 | 16126 16184 16189 16294 16296 16304 16519 73 198 263 | **T2b** |
| AMIS-R03 | 16008-255 | 16182C 16183C 16189C 16223T 16278T 16519C 73G 146C 153G | **X1'3** |
| AMIS-R04 | 15993-273 | 16256T 73G 263G | **P2** |

Table S2

Published HVS-I from Arabia, North Africa, East-Africa and Near East, collected for this study.

| Region/Population | Code | Number | Location | Reference(s) |
|---|---|---|---|---|
| **Arabia** | **ARA** | | | |
| Ta'izz | YE1 | 43 | Yemen | Černý et al. 2008 |
| Tihama | YE2 | 67 | Yemen | Černý et al. 2008 |
| Hajja | YE3 | 35 | Yemen | Černý et al. 2008 |
| Hadramawt | YE4 | 40 | Yemen | Černý et al. 2008 |
| Soqotra | YE5 | 65 | Yemen | Černý et al. 2009 |
| Saudi, Northern Region | SAN | 43 | Saudi Arabia | Abu-Amero et al. 2008 |
| Saudi, Western Region | SAW | 72 | Saudi Arabia | Abu-Amero et al. 2008 |
| **Arabia total** | | **365** | | |
| **Near East** | **NEE** | | | |
| Iraqi 1 | IR1 | 52 | Iraq | Al-Zahery et al. 2003 |
| Iraqi 2 | IR2 | 182 | Iraq | Al-Zahery et al. 2012 |
| Kurds | KUR | 50 | Eastern Turkey | Richards et al. 2000 |
| Syrians | SYR | 68 | Syria | Richards et al. 2000 |
| Turks | TUR | 131 | Turkey | Richards et al. 2000 |
| Jordanians | JOR | 145 | Jordan | Gonzalez et al. 2008 |
| **Near East total** | | **628** | | |
| **Northern Africa** | **NAF** | | | |
| Egypt Alexandria | EGA | 277 | Egypt | Saunier et al. 2009 |
| Egypt Lower | EGL | 54 | Egypt | Krings et al. 1999 |
| Egypt Upper 1 | EG1 | 33 | Egypt | Krings et al. 1999 |
| Egypt Upper 2 | EG2 | 58 | Egypt | Stevanovitch et al. 2003 |
| Nubians | NUB | 78 | Sudan Egypt | Krings et al. 1999 |
| Berbers Siwa | SIW | 78 | Egypt | Coudray et al. 2009 |
| Libyans | LIB | 269 | Libya | Fadhlaoui-Zid et al 2011 |
| Berbers Kesra | BKE | 47 | Tunisia | Cherni et al. 2005 |
| Arabs Zriba | AZR | 50 | Tunisia | Cherni et al. 2005 |
| Berbers Sened | BSE | 53 | Tunisia | Fadhlaoui-Zid et al. 2004 |
| Berbers Matmata | BMA | 49 | Tunisia | Fadhlaoui-Zid et al. 2004 |
| Berbers Chenini | BCH | 53 | Tunisia | Fadhlaoui-Zid et al. 2004 |
| Algerians | ALG | 240 | Algeria | Bekada et al. 2013 |
| Berbers Asni | BAS | 53 | Morocco | Coudray et al. 2009 |
| Berbers Bouhria | BBO | 70 | Morocco | Coudray et al. 2009 |
| Berbers Figuig | BFI | 94 | Morocco | Coudray et al. 2009 |
| Arabs North Tunisia | ANT | 64 | Tunisia | Turchi et al. 2009 |
| Arabs North Morocco | ANM | 56 | Morocco | Turchi et al. 2009 |
| **Northern Africa total** | | **1676** | | |
| **Eastern Africa** | **EAF** | | | |
| Dinka | DIN | 44 | Southern Sudan | Krings et al. 1999 |
| Sudanese | SUD | 102 | Sudan | Soares et al. 2012 |
| Ethiopians | ET1 | 77 | Ethiopia | Soares et al. 2012 |
| Amhara 2 | ET2 | 120 | Ethiopia | Kivisild et al. 2004 |
| Tigrais | ETE | 53 | Ethiopia Eritrea | Kivisild et al. 2004 |
| Gurage | ET3 | 21 | Ethiopia | Kivisild et al. 2004 |

| | | | | |
|---|---|---|---|---|
| Kenyans (Nairobi) | KE1 | 100 | Kenya | Brandstätter et al. 2004 |
| Dawro-Konta | ET4 | 137 | Ethiopia | Boattini et al. 2013 |
| Ongota | ET5 | 19 | Ethiopia | Boattini et al. 2013 |
| Hamer | ET6 | 11 | Ethiopia | Boattini et al. 2013 |
| Rendille | KE2 | 17 | Kenya | Boattini et al. 2013 |
| Elmolo | KE3 | 52 | Kenya | Boattini et al. 2013 |
| Luo | KE4 | 49 | Kenya | Boattini et al. 2013 |
| Maasai | KE5 | 81 | Kenya | Boattini et al. 2013 |
| Samburu | KE6 | 35 | Kenya | Boattini et al. 2013 |
| Turkana 3 | KE7 | 51 | Kenya | Boattini et al. 2013 |
| **Eastern Africa total** | | **969** | | |
| **Western Africa** | **WAF** | | | |
| Benin | BEN | 193 | Benin | Primativo et al. 2017 |
| **Eastern Africa total** | | **193** | | |
| **Total** | | **3831** | | |
